# Further Investigation of Mitochondrial Biogenesis and Gene Expression of Key Regulators in Ascites- Susceptible and Ascites-Resistant Broiler Research Lines

**DOI:** 10.1101/429431

**Authors:** Khaloud Al-Zahrani, Timothy Licknack, Destiny L. Watson, Nicholas B. Anthony, Douglas D. Rhoads

**Affiliations:** Program in Cell and Molecular Biology, University of Arkansas, Fayetteville, AR USA; Department of Biological Sciences, University of Arkansas, Fayetteville, AR USA; Department of Poultry Science, University of Arkansas, Fayetteville, AR USA

**Keywords:** Broiler, Hypertension, Gene expression, Mitochondria

## Abstract

We have extended our previous survey of the association of mitochondrial prevalence in particular tissues with ascites susceptibility in broilers. We previously reported that in breast muscle of 22 week old susceptible line male birds had significantly higher mtDNA copy number relative to nuclear copy number (mtDNA/nucDNA), compared to resistant line male birds. The higher copy number correlated with higher expression of *PPARGC1A* mRNA gene. Ascites is a significant metabolic disease associated with fast-growing meat-type chickens (broilers) and is a terminal result of pulmonary hypertension syndrome. We now report the mtDNA/nucDNA ratio in lung, liver, heart, thigh, and breast of both genders at 3, and 20 weeks old. At 3 weeks the mtDNA/nucDNA ratio is significantly higher in lung, breast, and thigh for susceptible line males compared to the resistant line males. Conversely, we see the opposite for lung and breast in females. At 20 weeks of age the differences between males from the two lines is lost for lung, and thigh. Although there is a significant reduction in the mtDNA/nucDNA ratio of breast from 3 weeks to 20 weeks in the susceptible line males, the susceptible males remain higher than resistant line males for this specific tissue. We assessed relative expression of five genes known to regulate mitochondrial biogenesis for lung, thigh and breast muscle from males and females of both lines with no consistent pattern to explain the marked gender and line differences for these tissues. Our results indicate clear sex differences in mitochondrial biogenesis establishing a strong association between the mtDNA quantity in a tissue-specific manner and correlated with ascites-phenotype. We propose that mtDNA/nucDNA levels could serve as a potential predictive marker in breeding programs to reduce ascites.

## Introduction

Ascites, Pulmonary hypertension syndrome PHS, or ‘water belly” is a cardiovascular, metabolic disease affecting fast-growing broilers. Ascites is a complex problem resulting from many interacting factors such as genetics, environment and management, but also occurs in normal conditions as a response to high metabolic rate [1–6]. The high metabolic oxygen requirement of rapid growth, combined with insufficient capacity of the pulmonary capillaries appears to be the most important cause of ascites incidence in modern broilers [7,8]. Inadequate oxygen levels trigger a series of events, including peripheral vasodilation, increased cardiac output, increased pulmonary arterial pressure, right ventricular hypertrophy (RVH; elevated right ventricular to total ventricular ratios- RV: TV), and ultimitly accumulation of fluid in the abdominal cavity and pericardium [5, 8, 9–11]. Advances in management practices, rearing programs, and improved selection techniques have decreased ascites incidence in modern broilers. However, ascites syndrome remains an economic concern throughout the world, causing estimated losses of $100 million annually in the US [12, 1993; Rossi personal communication, 2004; Cooper personal communication, 2018]. The etiology of ascites in poultry has been classified into three categories: 1) mainly pulmonary hypertension, 2) various cardiac pathologies, and 3) cellular damage caused by reactive oxygen species ROS [13]. Mitochondria are the powerhouses of the eukaryotic cell and are the major contributor to oxidative stress through the generation of reactive oxygen species (ROS). Mitochondria are the primary oxygen consumer for energy production to sustain rapid growth in broilers [14–16]. Mitochondria are known to be involved in the regulation of several fundamental cellular processes, including metabolism, apoptosis, intracellular signaling, and energy production in the form of ATP via the oxidative phosphorylation. Mitochondrial biogenesis can be defined as the process of growth and division of pre-existing mitochondria to increase ATP production in response to growing demand for energy or stress conditions [17]. During times of environmental stress (e.g., hypoxia, cold temperature, etc.), ROS levels can increase dramatically which may result in significant damage to cell structures notably the mitochondria [18]. Ascites can be induced at early ages by several methods such as altering the environment’s temperature [19, 20], air quality [21], and altitude

[22]. Researchers at the University of Arkansas established divergently selected ascites experimental lines derived from a former full pedigreed elite line beginning in the 1990s through sibling-selection based on a hypobaric challenge [11, 23]. The lines are the ascites resistant (RES) line, ascites susceptible (SUS) line, and a relaxed (REL) unselected line.

Previously we reported that for a small sample set of breast muscle at 22 weeks of age for RES and SUS males, the samples from SUS males had approximately twice the ratio of mitochondrial DNA (mtDNA) to nuclear DNA (nucDNA), and that this difference correlated with a difference in the level of expression of *PPARGC1A*[24]. We have further investigated this apparent difference and extended our analysis to genders, multiple tissues, and additional developmental stages. We also assessed the relative expression of five genes known to regulate mitochondrial biogenesis only for those tissues that demonstrated significant sex-differences in mtDNA copy number. The results indicate a likely correlation between mtDNA/nucDNA ratios and ascites phenotype for particular tissues.

## Materials and methods

### Birds stocks

All animal procedures were approved by the University of Arkansas Institutional Animal Care and Use Committee (under protocol 12039 and 15040). Birds used in this study represent the ascites-resistant (RES), the ascites-susceptible (SUS), and the relaxed unselected (REL) lines at generation 21 [23].

### Tissue collection

Heart, lung, muscle iliotibialis (thigh), pectoralis major (breast), and liver, were collected from SUS and RES experimental lines. At three weeks of age, five male and female birds from each experimental line were randomly selected, euthanized by cervical dislocation, and samples were collected and immediately stored in RNAlater^TM^ (Sigma Aldrich, St. Louis, MO). At 20 weeks of age we collected lung, thigh, and breast from five males of SUS and RES lines. For the REL line we collected breast tissue from 12 males at 3 and 20 weeks of age.

### DNA isolation

Tissue samples were homogenized in 1 ml lysis buffer (10 mM TrisCl, 1 mM Na2EDTA pH 7.5) using a Bullet Blender homogenizer (Next Advance, Inc., Averill Park, NY) and overnight digested with 100 µg/ml pronase at 37°C. SDS was added to, then successively extracted by phenol:chloroform:isoamyl alcohol (25:24:1) and chloroform:isoamyl alcohol (24:1), followed by ethanol precipitation of DNA. DNAs were dissolved in 10 mM TrisCl 0.1 mM EDTA pH 7.5. DNA quantity was assessed by fluorimetry with Hoechst 33258 (GLOMAX Multi Jr, Promega Corp., Madison, WI) and purity (A260/280) by spectrophotometry (NanoVue, GE Healthcare Bio-Sciences, MA, USA).

### RT-qPCR for mitochondrial biogenesis

Mitochondrial DNA content was measured by quantitative, real time PCR (qPCR) in 96 well format using a CFX96 Thermal Cycler (Bio-Rad Laboratories, Inc., Hercules, California, USA). The mitochondrial target was the gene for mt-tRNA^ARG^, with the nuclear target a region of 5-Hydroxytryptamine receptor 2B (HTR2B). Specific primers (Table 1) were designed using Primer3 software (version 0.4.0; http://bioinfo.ut.ee/primer3–0.4.0/primer3/) and synthesized by Integrated DNA Technologies (Coralville, IA USA). Reactions (20 μl) were run in triplicate and consisted of 1X Taq Buffer (50 mM Tris-Cl, pH 8.3, 1 mM MgCl_2_, 30 μg/mL BSA), 1X EvaGreen dye (Biotium Inc., Hayward, California, USA), 0.25 mM MgCl_2_, 0.2 mM dNTP, 0.5μM each of the specific forward and reverse primers, 4 U of Taq polymerase, 2 μL of DNA (50–100ng). The cycling protocol was an intial soak at 90°C for 30 s, followed by 40-cycles of 30 s at 95 °C, 15 s at 60°C and 30 s at 72 °C followed by a plate read. Ct values from the exponential phase of the PCR were exported directly into Microsoft EXCEL worksheets for analysis. The ∆Ct of mtDNA relative to the nucDNA reference SUS samples were converted to ΔΔCt values calibrated based on the ∆Ct of RES samples [25]. The fold changes relative to the calibrator (RES line) was estimated as 2^(-ΔΔCt)^.

**Table 1.**
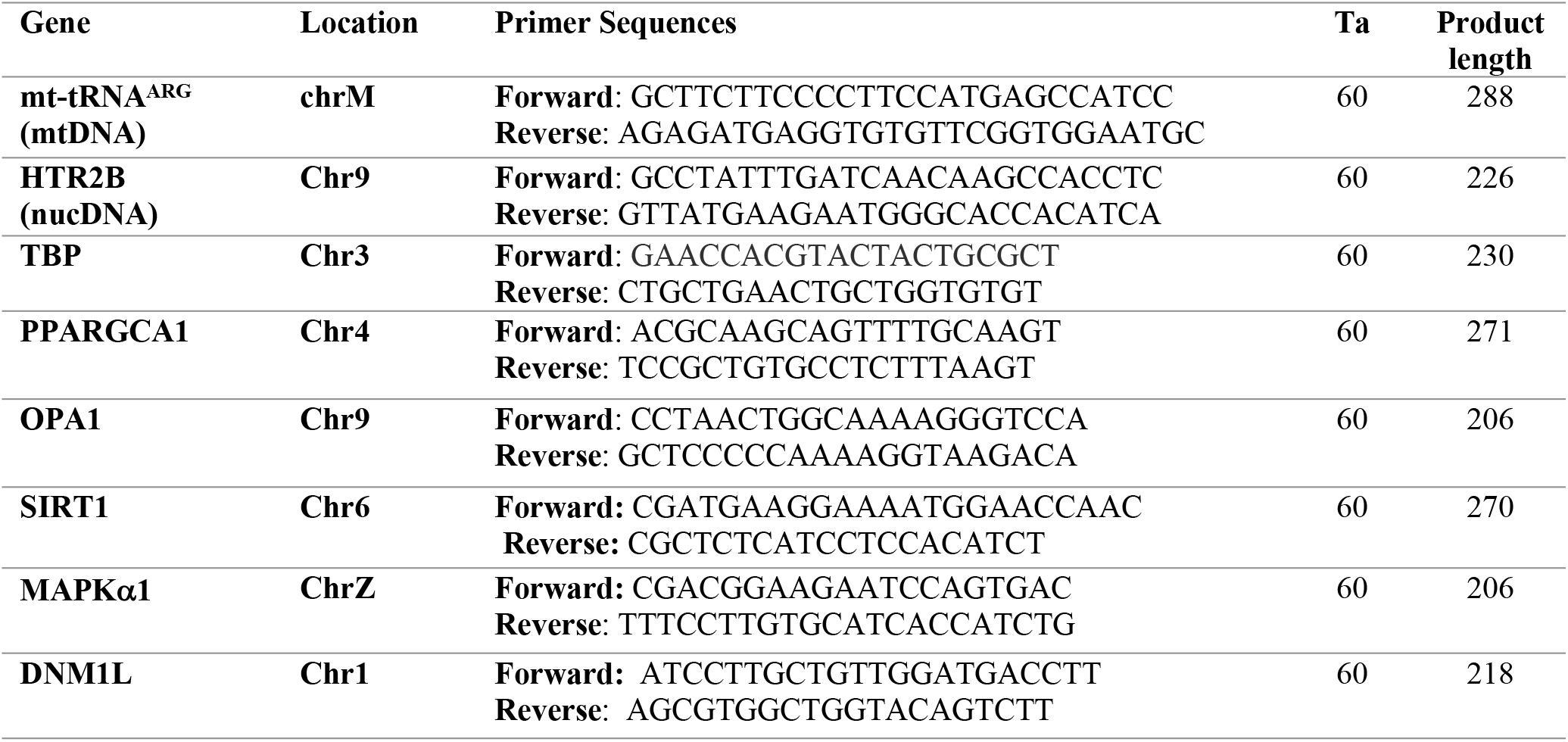
Sequences of primer pairs used for RT-qPCR analysis of chicken target and reference genes. For each gene the primer sequence for forward (F) and reverse (R) are listed (5’-3’), genomic location, the annealing temperature in °C used (Ta), the amplicon product length (bp). All primer sequences were synthesized by Integrated DNA Technologies (IDT, Coralville, IA).

### RNA isolation and gene expression analyses

Total RNA was isolated from lung, thigh, and breast tissues using TRIZOL reagent (Ambion, Thermo Fisher Scientific) according to the manufacturer’s instruction. The extracted RNA was assessed for quantity and purity (A260/280) using NanoVue spectrophotometry (GE Healthcare Bio-Sciences, MA, USA). RNA integrity was evaluated by electrophoresis in 1.5% agarose gel in 0.5×TBE buffer (50 mM Tris, 1 mM Na_2_EDTA, and 25 mM Borate, pH 8.3), stained by 0.5 μg/ml ethidium bromide. Samples that did not show 3 strong and distinct bands (28S, 18S, and 5S rRNA) were discarded. Gene expression for *PPARGC1A, AMPαK, OPA1, SIRT1*, and *DNM1L* was performed using a two-step RT-qPCR method. RNA (up to 5μg) was combined with 2 μM CT_23_V, and 0.5 mM dNTP and denatured at 70°C for 5 mins, then added to a mastermix consisting of 1X First Strand buffer (Invitrogen), 5 mM MgCl_2_, 1 mM DTT, 20 U RNasin (Promega Corp, Madison, WI USA), and 200 U MMLV reverse transcriptase (Promega Corp) in a final volume of 20 μl. The reaction was incubated at 42°C for 60 minutes and then inactivated at 85°C for 5 minutes. Chicken TATA-binding protein (*TBP*) was used as the reference gene [26]. Primers (Table 1) for each gene were designed to span an intron using Primer3 software and synthesized by Integrated DNA Technologies. Second step qPCR were performed in a 20μl volume were as above for qPCR except as target 2 μL of cDNA (50–100ng). The PCR cycling was initial denaturation at 90°C for 3 mins, 10 cycles of 90°C for 15s, 60 °C for 15s, 72 °C for 1 min, followed by another 30 cycles of 90°C for 15s, 60 °C for 15s, melt curve 70°C to 90°C, finally 72 °C for 1 min with plate read. Ct values were analyzed as described above. Relative gene expression was calculated using the 2^(-ΔΔCt)^ method [25] with both biological and technical replicates, and normalized to TBP as the reference gene.

### Statistical analysis

Data are presented as means ± SEM. All statistical computations were performed using EXCEL, and significant difference between lines and gender means were assessed by the Student’s t-test. Probability level of P ≤ 0.05 was considered statistically significant.

## Results

Previously we evaluated the mitochondrial biogenesis and *PPARGC1A* mRNA gene expression in male broiler chickens at 22 weeks of age [24]. The analyses compared two experimental lines produced through divergent selection for ascites phenotype; the ascites-susceptible (SUS) and ascites resistant (RES) broiler lines. The comparison was based on five males from each line and the evaluation was for right ventricle and breast. Results showed that birds from SUS had significantly higher mtDNA copy number (P = 0.038) and PPARGC1A RNA gene (P = 0.033) in breast muscle; with no difference in right ventricle. Thus, we suggested that mitochondrial biogenesis and *PPARGC1A* mRNA gene expression differ between male boilers from RES and SUS lines in a tissue-specific manner. The present report extends our previous analyses to additional muscles and other critical tissues at additional ages and for both genders.

From each line, five birds of both sexes were sampled for right ventricle, breast, thigh, lung, and liver at 3 weeks of age. The mtDNA/nucDNA ratio was estimated by qPCR of mt-tRNA^ARG^

(mtDNA) and a single copy region of HTR2B (nucDNA). A higher mtDNA/nucDNA ratio was observed in lung (Fig 1.C), thigh (Fig 1.D), and breast (Fig 1.E) tissues of SUS line relative to the RES line in males. The breast tissue of SUS line males contained 4 times higher levels (P=0.048) of mtDNA copy number. The lung of SUS line males was 64 times higher (P=0.01) and the thigh was 16 times higher (P= 0.03). No differences were detected in mtDNA/nucDNA ratio between the males from the two lines for right ventricle (Fig 1.A), and liver (Fig 1.B). Although the right ventricle of SUS line males was higher than RES line males, the difference was not statistically significant (P = 0.08). Inspection of the mtDNA/nucDNA ratios across tissues for males from each line revealed that the RES line males were comparable (around 1000) for right ventricle, thigh, liver and breast, but only around 100 for lung. The SUS males were much more variable ranging from 100,000 for thigh and lung, to 5,000 to 10,000 for liver, right ventricle, and breast.

**Fig 1:**
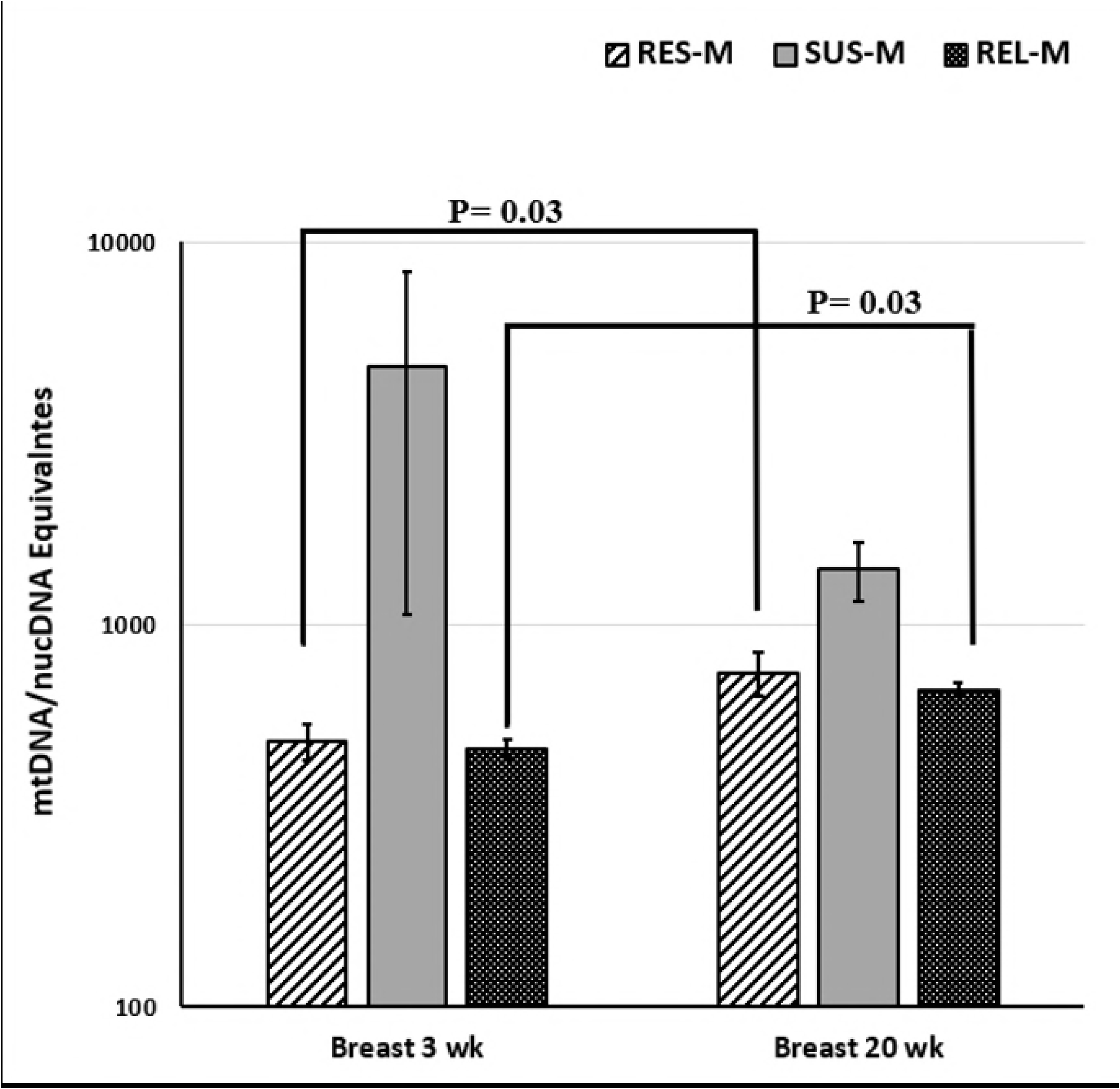
Mitochondrial Biogenesis in several tissues from ascites research lines. (A) and (B) Mean mtDNA relative to nucDNA in heart, and liver of the SUS and RES lines of both sexes at 3 weeks old. (C), (D), and (E) Mean mtDNA relative to nucDNA in lung, thigh, and breast muscle of the SUS and RES lines of both sexes at 3 and 20 weeks old (n=5 for each group). Error bars are SEM and P values determined by one-tailed t-test, P <0.05.

In contrast to the males, the mtDNA/nucDNA ratio at 3 weeks of age for SUS line females were lower than RES line females for lung (Fig 1.C) and breast (Fig 1. E). The breast ratio for SUS line females was half that of the RES females (P=0.03), while for lung the SUS line was 0.008x the value for the RES females (P= 0.004). No differences in mtDNA/nucDNA ratio were observed between the females for the two lines for liver (Fig 1.B), right ventricle (Fig 1.A), and thigh (Fig 1.D). Although the liver, and right ventricle of SUS line females was lower than RES line females, the difference was not statistically significant, and the RES line female values for liver were more variable. Examination of the mtDNA/nucDNA ratios across tissues for females for both lines revealed that the SUS line females were comparable (around 1000) for right ventricle, thigh, lung, and breast, and around 10,000 for liver. Unlike for males the RES female samples showed the greatest tissue variation. RES females’ ratios ranged from 100,000 for liver and lung, to 10,000 for right ventricle, and 1000 for thigh and breast.

Comparison of mtDNA copy number between genders within each line at 3 weeks of age shows significant differences for some tissues. Females from the RES line had higher mtDNA copy number than males from the RES line for lung (P=0.001) and breast (P=0.006). In contrast, males from the SUS line had a relatively higher mtDNA copy number than SUS line females for lung (P= 0.05) and thigh (P=0.03). In this study, lung tissue demonstrated the most significant mtDNA/nucDNA ratio differences in respect to both gender and line.

Since only lung, thigh and breast showed differences at 3 weeks of age, we examined mtDNA/nucDNA ratios for those same tissues at 20 weeks of age. We restricted our investigation to males since ascites mortality is consistently higher for males than for females in our research lines. This is also consistent with reports from other researchers on commercial broilers [27–30]. Five males of both lines from the same generation were assessed for ontological changes in mtDNA copy number. As shown in Fig 1.C, and 1.D, we observed a decrease in mtDNA/nucDNA ratio in 20 week SUS line males compared to 3 week SUS line males for lung (P= 0.019) and thigh (P= 0.045). The reduction at 20 weeks of age results in the SUS and RES line males have comparable levels of mtDNA for lung and thigh. However, as for 3 weeks of age we continue to see a higher ratio of mtDNA/nucDNA in the breast muscle for the SUS line males compared to the RES line males (Fig 1.E). The difference between the lines for the breast decreases from 6-fold at 3 weeks of age to approximately 2-fold at 20 weeks of age but remains statistically different between the lines (P=0.02). Furthermore, consistent with our finding at a younger age, the mtDNA/nucDNA ratio in breast muscle of 20 weeks old of SUS line males was 2 times higher (P= 0.02) compared to RES line. No differences between young and old birds in mtDNA/nucDNA ratio were detected for breast tissues of SUS line males (P=0.3) indicating the consistent elevation of mtDNA/nucDNA ratio. However, we observed an increase in mtDNA/nucDNA ratio in breast tissues (P=0.03) between young and old birds of the of RES line. Thus at 20 weeks of age we see the SUS line male mtDNA copy number reduced to comparable levels as the RES line males for lung and thigh but the breast mtDNA copy number remains elevated in the SUS line compared to the RES line.

Since the REL line represents the founder population for our SUS and RES experimental lines, we decided to examine the mtDNA/nucDNA ratio in breast muscle of male birds from the REL line at 3 and 20 weeks of age. As shown in Fig 2, we observed an increase in mtDNA/nucDNA ratio in 20 week REL line males compared to 3 week REL line males (P=0.03). In this study, both REL and RES line male birds had the same mtDNA/nucDNA ratio at both 3 and 20 weeks of age. Unlike the SUS male birds that have always higher mtDNA/nucDNA ratio at both ages than the RES and REL males.

**Fig 2.**
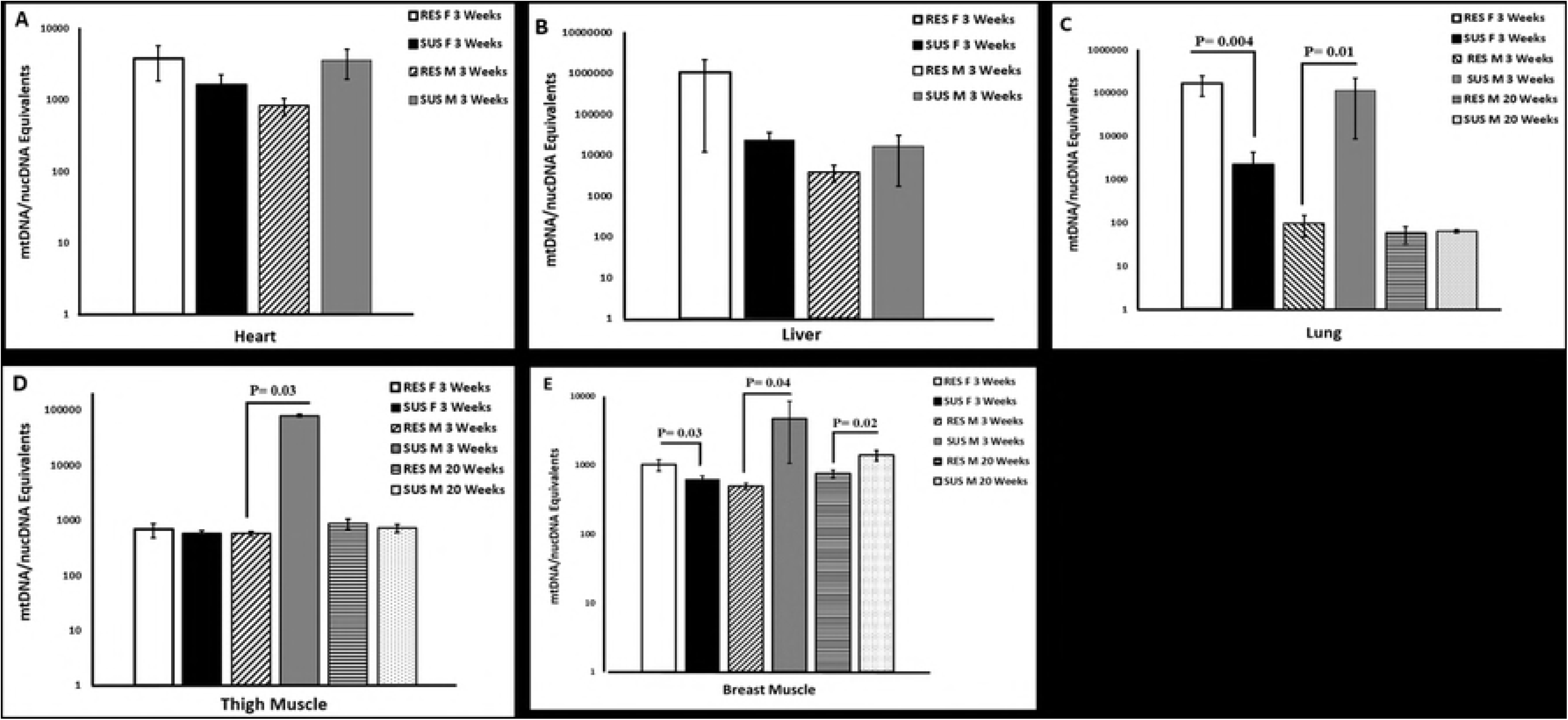
Mitochondrial Biogenesis in breast muscle of 3 and 20 weeks old male birds. mtDNA relative to nucDNA in breast tissues of males from RES, SUS, and REL experimental lines at 3 weeks old, and 20 weeks old (n=5 for SUS and RES birds and n=12 for REL line birds). Error bars are SEM and P values determined by one-tailed t-test, P <0.05.

A number of genes have been associated with regulation of mitochondrial biogenesis. We selected five of these genes to examine their expression levels using RT-qPCR for the tissues showing the greatest differences for gender or line. The five genes were: AMP-activated protein kinase α1 (*AMPKα1)*, peroxisome proliferator-activated receptor gamma co-activator 1 alpha (*PPARGC1A)*, Sirtuin 1 (*SIRT1)*, optic atrophy 1 (*OPA1), and* Dynamin-1 like (*DNM1L)*. The expression of these genes were assessed in lung, thigh, and breast of both lines and genders at 3 weeks of age, and breast and lung for males at 20 weeks of age. In all cases the relative expression was determined and calibrated against the expression in the RES line.

In males at 3 weeks of age, expression of all five genes were reduced in all three tissues (Table 2), with the reduction being statistically significant for *AMPKα1, OPA1, and DNM1L* in lung and breast ranging to half the expression in SUS relative to RES. There were no differences in the level of expression PPARGC1A and SIRT1 genes in lung and breast between the two lines at this age. In thigh, there were no differences in expression levels for any of these five genes. Interestingly, an increase in the level of expression of *PPARGC1A, SIRT1*, and *OPA1* genes in breast of SUS males at 20 weeks of age relative to RES males was observed. No differences in the expression of DNM1L gene in breast while the expression of *AMPKα1* gene remained low. Inthe lungs of SUS males at 20 weeks of age, a reduction in the expression of all genes was observed relative to RES males (Table 3), with the reduction being significant for *AMPKα1* (P=0.016), and *OPA1*(P= 0.0009). No significant differences in *PPARGC1A* expression was observed.

**Table 2:**
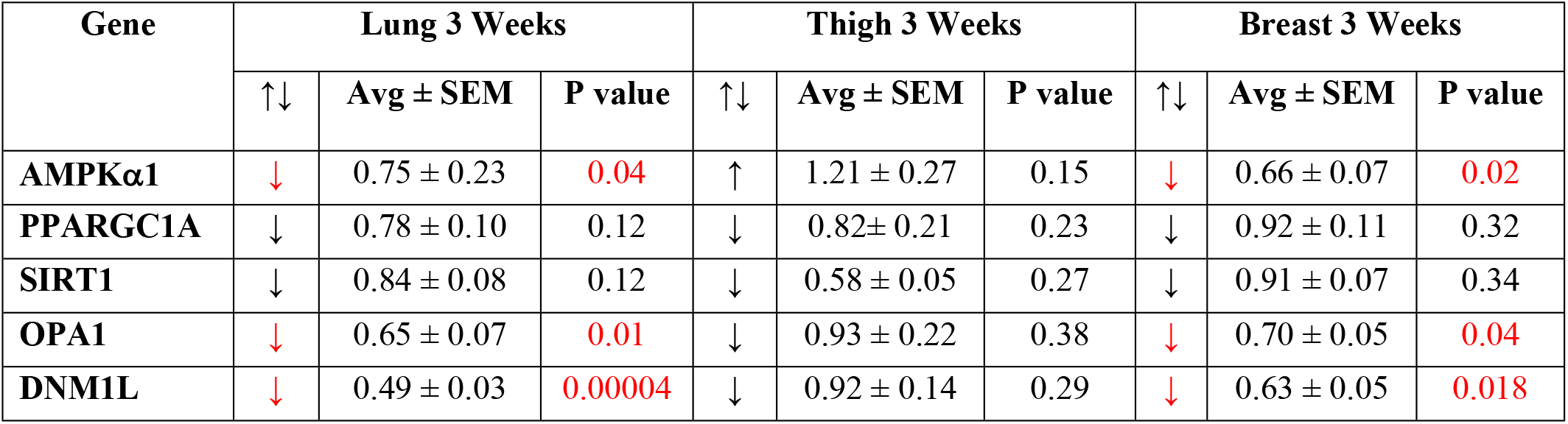
Relative gene expression in lung, thigh, breast muscles of 3 weeks old male birds from SUS and RES lines divergently selected for ascites phenotype. ↑↓ indicates the direction of the difference for the SUS line relative to the RES line. Avg ± SEM is the average ± standard error of the mean for the n-fold change (2^-ΔΔCt) for the SUS line relative to the calibrator, RES line, from five birds (n= 5) run in triplicate. Statistically different results were determined using Student’s *t* test for unpaired samples.

| Gene | Lung 3 Weeks |  |  | Thigh 3 Weeks |  |  | Breast 3 Weeks |  |  |
| --- | --- | --- | --- | --- | --- | --- | --- | --- | --- |
| | $\uparrow\downarrow$ | Avg ± SEM | P value | $\uparrow\downarrow$ | Avg ± SEM | P value | $\uparrow\downarrow$ | Avg ± SEM | P value |
| **AMPKα1** | ↓ | 0.75 ± 0.23 | **0.04** | ↑ | 1.21 ± 0.27 | 0.15 | ↓ | 0.66 ± 0.07 | **0.02** |
| **PPARGC1A** | ↓ | 0.78 ± 0.10 | 0.12 | ↓ | 0.82 ± 0.21 | 0.23 | ↓ | 0.92 ± 0.11 | 0.32 |
| **SIRT1** | ↓ | 0.84 ± 0.08 | 0.12 | ↓ | 0.58 ± 0.05 | 0.27 | ↓ | 0.91 ± 0.07 | 0.34 |
| **OPA1** | ↓ | 0.65 ± 0.07 | **0.01** | ↓ | 0.93 ± 0.22 | 0.38 | ↓ | 0.70 ± 0.05 | **0.04** |
| **DNM1L** | ↓ | 0.49 ± 0.03 | **0.00004** | ↓ | 0.92 ± 0.14 | 0.29 | ↓ | 0.63 ± 0.05 | **0.018** |

**Table 3:** Relative gene expression in lung, and breast muscles of 20 weeks old male birds from SUS and RES lines divergently selected for ascites phenotype. Column headers and data representations are as described for Table 2. NC is no significant change.

| Gene | Lung 20 Weeks |  |  | Breast 20 Weeks |  |  |
| --- | --- | --- | --- | --- | --- | --- |
|  | ↑↓ | Avg ± SEM | P value | ↑↓ | Avg ± SEM | P value |
| **AMPKα1** | ↓ | 0.15 ± 0.13 | 0.01 | ↓ | 0.83 ± 0.29 | 0.02 |
| **PPARGC1A** | NC | 1.01 ± 2.05 | 0.16 | ↑ | 1.97 ± 0.09 | 0.000003 |
| **SIRT1** | ↓ | 0.51 ± 0.08 | 0.10 | ↑ | 1.43 ± 0.11 | 0.004 |
| **OPA1** | ↓ | 0.62 ± 0.05 | 0.0009 | ↑ | 1.18 ± 0.07 | 0.03 |
| **DNM1L** | ↓ | 0.81 ± 0.08 | 0.11 | ↓ | 0.92 ± 0.23 | 0.19 |

In females at 3 weeks of age, the expression of *AMPKα1* gene in lung was reduced (0.5x; P=0.05) in SUS line compared to RES line, while *SIRT1* mRNA expression increased by approximately 29% (Table 4). For thigh, only *OPA1* and *DNM1L* were reduced to 80% and 60%, respectively, in SUS vs RES females. In breast, none of the five genes were found to differ between the lines. No gene expression analyses were performed for females at 20 weeks of age.

**Table 4:** Relative gene expression in lung, thigh, breast muscles of 3 weeks old female birds from SUS and RES lines divergently selected for ascites phenotype. Column headers and data representations are as described for Table 2. NC is no significant change.

| Gene | Lung 3 Weeks |  |  | Thigh 3 Weeks |  |  | Breast 3 Weeks |  |  |
| --- | --- | --- | --- | --- | --- | --- | --- | --- | --- |
|  | ↑↓ | Avg ± SEM | P value | ↑↓ | Avg ± SEM | P value | ↑↓ | Avg ± SEM | P value |
| **AMPKα1** | ↓ | 0.43 ± 0.50 | 0.05 | ↓ | 0.87 ± 0.28 | 0.19 | NC | 1.03 ± 0.24 | 0.47 |
| **PPARGC1A** | ↓ | 0.48 ± 0.18 | 0.12 | NC | 1.01 ± 0.13 | 0.49 | ↓ | 0.48 ± 0.16 | 0.08 |
| **SIRT1** | ↑ | 1.29 ± 0.12 | 0.02 | ↓ | 0.98 ± 0.07 | 0.41 | ↓ | 0.92 ± 0.09 | 0.28 |
| **OPA1** | ↑ | 1.16 ± 0.17 | 0.22 | ↓ | 0.80 ± 0.07 | 0.03 | ↓ | 0.86 ± 0.05 | 0.15 |
| **DNM1L** | ↓ | 0.60 ± 0.19 | 0.07 | ↓ | 0.60 ± 0.08 | 0.01 | ↓ | 0.79 ± 0.18 | 0.40 |

None of these key mitobiogenesis regulators appeared to correlate with the differences in mtDNA/nucDNA ratios we observed for both genders between the two lines. However, in males, we observed a reduction in the mtDNA copy number for SUS males from 3 weeks of age to 20 weeks of age in lung, thigh, and breast, although the levels remained relatively higher in breast of the SUS males than the RES males. Consistent with that difference at 3 weeks of age we saw a decreased expression for *OPA1, and DNM1L* genes in lung and breast SUS males relative to RES males whereas at 20 weeks of age we observed a higher expression for PPARGC1A, SIRT1, and *OPA1* in only breast of SUS males relative to RES males. Additionally, *AMPKα1* gene was always expressed at lower levels in the breast and lung tissues of SUS males compared to RES males at both ages. In general, we found no consistent gene expression pattern to explain the marked gender and line differences in mtDNA copy number for these tissues.

## Discussion

Mitochondrial dysfunction is well documented in a wide array of diseases and conditions, such as Alzheimer’s disease, cancer, and aging [31–33]. Mitochondria are central to ATP synthesis, heat production, radical oxygen species (ROS) generation, fatty acid and steroid metabolism, cell proliferation, and apoptosis [17; 34]. Alterations in mtDNA sequence or copy number may contribute to mitochondrial dysfunction [35]. Thus, it is likely that imbalances within the cell concerning mitochondria-centered metabolic pathways may contribute to ascites syndrome. Our observations indicate that variations in mtDNA copy number could be an important component in the pathoetiology of ascites syndrome in broilers. Using different tissues, we have demonstrated that mtDNA copy number can be an important biomarker during early developmental age for ascites syndrome susceptibility. Our results showed sizable tissue-specific, and gender differences in the mtDNA/nucDNA ratio at early ages of broilers. The possible existence of gender-specific differences in energy metabolism for particular tissues might be a consequence of interplay between maternally inherited mitochondria and sex chromosomes or differences in endocrine responses. In males, mtDNA/nucDNA ratio was significantly higher in lung, thigh, and breast tissues from SUS line males at 3 weeks of age in comparison with RES line males. Conversely, mtDNA levels were significantly lower in lung and breast tissues of SUS line females as compared to RES line females. The gender differences may impact ascites phenotype considering that males are documented to have higher ascites mortality than females [27–30] The observed elevation in the amount of mtDNA in lung, thigh and breast muscle of SUS line males might be attributable to a compensatory response to the decline in the respiratory function of mitochondria or a response to other metabolic regulatory processes. An alternate explanation is needed for the reduced mtDNA copy number in lung and breast muscle of SUS line females. One possible explanation is that differences in mtDNA content of different sexes can be attributed to imbalances in oxidative stress due to higher female estrogen levels. Previous work found that oxidative damage to mtDNA is 4-fold higher in males than in females [36,37]. The lower oxidative damage in females may be attributable to the protective effect of estrogens by upregulating the expression of antioxidant enzymes in mitochondria via intracellular signaling pathways, thus decreasing oxidative damage and increasing antioxidants defenses [36,37]. Moreover, fundamental sex differences in metabolism under stressful conditions have long been observed in several organisms and may also be influenced by intrinsic differences in genomic maintenance [38]. Absent from our analysis is any determination of whether the differences in mtDNA content is associated with functional or non-functional (defective) mitochondria. Future work could involve fluorescent detection systems for visualizing mitochondria in SUS vs RES tissues to assess relative mitochondrial abundance and functional state.

Bottje and Wideman hypothesized that mitochondrial dysfunction contributes to systemic hypoxia that leads to ascites in broilers [9]. They reported that mitochondrial function is defective in a variety of tissues (lung, liver, heart, and skeletal muscles) in male broilers with ascites where oxygen utilization is less efficient than in male broilers without ascites [4, 15, 16,39]. They assessed the mitochondria function for both the respiratory control ratio (RCR); for electron transport chain coupling, and for the adenosine diphosphate to oxygen ratio (ADP:O); for oxidative phosphorylation. A decline in RCR and ADP:O ratio was detected in ascites mitochondria relative to the non-ascites control. This may indicate functional impairment of mitochondrial oxidative phosphorylation and less efficient utilization of oxygen than in control. On the other hand, more efficient oxidative phosphorylation and lower oxidative stress were observed in mitochondria obtained from broilers selected for genetic resistance to ascites. Accumulation of hydrogen peroxide was observed in heart and skeletal mitochondria in broilers with ascites and of oxygen radical production in ascites liver and lung mitochondria. Therefore, there is no doubt that mitochondrial function is defective in broilers with ascites which leads to increased production of ROS. It is possible that the observed significantly lower mitochondrial biogenesis in male RES and REL lines is indicative of lower oxygen demand or ROS production. However, it is yet not clear if increased levels of ROS are a secondary effect of development of ascites or are associated with genetic susceptibility. Cisar et al. (2004) used immunoblots to quantify cardiac mitochondrial electron transport chain (ETC) protein levels in the RES and SUS lines under hypoxic challenge. ETC protein levels were similar in RES and SUS at ambient oxygen pressure but were significantly elevated only in RES under hypoxic conditions. Our data is for ambient oxygen levels only and based on mitochondrial DNA and not mitochondrial proteins sugessting the possible involvment of mitochondrial proteins with ascites phenotype [40].

Imbalance in mitochondrial biogenesis may only affect broilers at a young age when ascites is most likely to develop. Contrary to 3 weeks of age, at 20 weeks of age males from the RES and SUS lines showed no differences in mtDNA copy number for lung and thigh. One additional potentially confounding aspect is that the 20 week samples were from birds that had been feed restricted since 5 week post hatch. Despite this, the difference in breast mtDNA copy number was still higher for SUS males compared to RES males. Future investigations should examine females for a similar ontological shift in mitochondrial biogenesis, as well as assess the impact of feed restriction. Apparently, the consistent increase in mtDNA/nucDNA ratio between young and old birds of the two lines is restricted to breast muscle which may reflect increased energy demands or a compensatory amplification to overcome the loss of mitochondrial function or oxidative stress.

Examination of mtDNA/nucDNA ratio in breast muscle from the REL line of male birds at 3 and 20 weeks of age indicate a similar pattern as the RES line. This was surprising since our research lines, SUS and RES, were originally developed from the REL line. We expected to see a wider range of mtDNA abundance in the REL line reflecting a composite of the SUS and RES patterns. This may be the result of imbalance between the rate of biogenesis and clearance of dysfunctional or old mitochondria in SUS vs REL and RES males. Alternatively, this may be due to imbalance in mitochondrial-nuclear crosstalk is SUS vs REL and RES males. Our study strongly supports a potential decrease in the mitochondrial function with oxidative stress, yet overall mtDNA quantity increases by a feedback mechanism to compensate for general mitochondrial dysfunction and damage in ascites-SUS male birds. However, the detailed mechanism remains unclear.

We analyzed gene expression of some of the key regulators of the mitochondrial biogenesis in ascites- susceptible and ascites- resistance lines of both genders. *PPARGC1A* is the master regulator of mitochondrial biogenesis. This transcriptional coactivator coordinates the actions of several transcription factors that involved in the basic functions of the mitochondrion as well as its rate of biogenesis [41, 42]. No changes in the *PPARGC1A* expression were detected in lung, thigh, and breast tissues of both genders and lines at 3 weeks of age. However, consistent with the observed increased mitochondrial biogenesis in in breast tissue of the SUS males at 20 weeks old, the levels of PPARGC1A mRNA gene expression were almost 2-fold change higher relative to the RES birds. Probably, the increased activity of PPARGC1A in breast muscle during sexual maturity could play a role in enhancing mitochondrial respiratory capacity which attenuates the development of ascites in SUS males. However, it has yet to be determined whether the enhanced activity of *PPARGC1A* is attributable to its promotion of mitochondrial function or its effects on nonmitochondrial gene expression.

*AMPKα1* gene regulates intracellular energy metabolism in response to acute energy crises and is activated by an increase in AMP/ ATP ratio (energy depletion) and inhibited by the presence of glycogen. Thus, to maintain energy homoeostasis, *AMPKα1* switches on catabolic pathways that generate ATP, while switching off anabolic pathways that consume ATP [42, 43]. Interestingly, The *AMPKα1* gene activity was notably down regulated in lung, and breast tissues at 3 and 20 weeks of age of SUS line males in comparison with RES line. In females, *AMPKα1* gene was only downregulated in lung muscle of SUS line birds as compared to RES line at early age. Several studies indicate another important role of *AMPKα1* in the disposal of dysfunctional and damaged mitochondria, process known as autophagy [43]. Any impairment of the mitochondrial autophagy process is often accompanied by accumulation of dysfunctional or damaged mitochondria that leads to increases in mtDNA content. Therefore, it is possible that the observed *AMPKα1* downregulation in SUS line as compared with the RES line caused insufficient removal of the damaged mitochondria which may explain the increase mtDNA content in male birds of the SUS line as compared with the RES line.

*OPA1* gene plays an essential role in the inner mitochondrial fusion and maintenance of the mitochondrial network architecture, which is essential for mitochondrial activity and biogenesis. *DNM1L* is the master regulator of mitochondrial division in most eukaryotic organisms [44]. Remarkably, at 3 weeks of age, both *OPA1* and *DNM1L* mRNA expression were significantly decreased in lung, and breast tissue of SUS line males, and in thigh of SUS line females as compared with RES line. Downregulation of *DNM1L* and *OPA1* genes in these tissues at this early age may reduce the efficiency of mitochondrial autophagy and causes accumulation of dysfunctional mitochondria. Consequently, the mitochondria are not able to re-fuse with the mitochondrial network after fission leading to increase in fragmented mitochondria and mtDNA accumulation. Conversely, at 20 weeks old, the *OPA1* was found to be significantly upregulated in breast tissue of SUS line males as compared with RES line which may reflects the enhanced activity in the mitochondrial biogenesis and the quick clearance of damaged mitochondria in breast muscle as birds advance in age.

*SIRT1*, a metabolic sensor that belongs to the sirtuin (NAD+ –dependent deacetylases) family and its activity can increase when NAD^+^ levels are abundant, such as times of nutrient deprivation. SIRT1 stimulates mitochondrial biogenesis via deacetylation of a variety of proteins in response to metabolic stress [42, 45]. In our study, *SIRT1* was overexpressed in breast tissue of SUS line males at 20 weeks old and in lungs of SUS line females at 3 weeks of old compared to RES birds.

In summary, our findings indicate clear sex differences in mitochondrial biogenesis establishing a strong association between the mtDNA content and ascites-susceptibility and ascites-resistance in a tissue-specific manner. The mtDNA/nucDNA levels could serve as potential predictive markers to screen for ascites phenotype in birds at early developmental ages. Moreover, this study confirms that the consistent increase in the mtDNA/nucDNA ratio between young and old birds is only restricted to breast muscles. However, it is worth noting that mitochondrial biogenesis is tissue specific. This is because every type of cell and tissue has a specific transcriptional profile, and consequently unique features of metabolic pathways. Our study suggests the possible contribution of the lower expression of *AMPKα OPA1*, and *DNM1L* genes in mitochondrial biogenesis defects in male SUS birds which leads to increase in mtDNA content in some tissues at early ages. Furthermore, our data is consistent with a possible role of *PPARGC1A* in breast tissue of SUS line males in controlling ascites syndrome progression and improved regulation of mitochondrial biogenesis at older ages. Nevertheless, we have no clear evidence for what genes or regulators are driving the observed sizable sex-differences in mtDNA copy number at an early age. Despite our findings, the precise mechanism that explains the association between mtDNA copy number and ascites syndrome remains unknown. To address this further in the future, we need to test larger sample numbers, more tissues, and different populations/crosses. Our observations are based on a single experimental series and, although our results agreed with our previous data, we cannot wholly determine if this phenomenon is a cause or effect or limited to the tissues used in this study. Regardless of the limited number of replicates used, our study had sufficient statistical power to detect significant differences in mtDNA/nucDNA ratio and gene expression analysis. Future research should focus on finding mitochondrial biogenesis causal genetic regulators and exploring whether they are connected or unrelated to changes in the mtDNA.

## Acknowledgements

This project was supported by Agriculture and Food Research Initiative Competitive Grant no. 2014–06315 from the USDA National Institute of Food and Agriculture to DR and NA. KA was supported on a fellowship through the Saudi Arabia Cultural Mission.

